# Long term trend and short-term dynamics of a willow ptarmigan population

**DOI:** 10.1101/2023.05.03.539219

**Authors:** Tomas Willebrand, Eivind Flittie Kleiven, Maria Hörnell Willebrand, Rolf Brittas

## Abstract

Willow ptarmigan *Lagopus lagopus* is abundant in Arctic and tundra regions, but rapid climate warming has raised concerns about possible declines as has been observed in several bird species. In this study, we used a hierarchical state-space model to analyze data from a 139 km line transects in mid-Sweden over 48 years. Adult numbers and breeding success were analyzed separately, and we included covariates on vole abundance, numbers of snow-free days in autumn and spring, and the last day of frost in May. We assessed long-term trends in the adult population and estimated the effects of breeding success and weather variation on short-term changes. The estimates of adult density did not show any trend for the period 1976 to 2023, and the dynamics were characterized by a strong direct negative density dependence indicating a stationary process. A number of possible mechanisms have been suggested for how a warmer climate affects willow ptarmigan population dynamics, but our results do not support the hypothesis that lack of snow in autumn and spring increases the vulnerability of willow ptarmigan to predation and leads to population decline. Breeding success is an important driver of changes in, but independent of, adult density. In addition to predation, we propose that climate conditions and emerging vegetation during egg formation and laying is important. We suggest that our results can be explained by a diverse predator assemblage that makes it difficult for the population to escape top-down control, resulting in short-term fluctuations at lower densities.

## Introduction

Negative trends and population declines have been reported for several bird species in the northern hemisphere, and climate change has been identified as the most important process behind the declines (Smith et al., 2020). Willow ptarmigan *Lagopus lagopus* is an abundant herbivore found in the Arctic and tundra regions (Storch, 2007). It has an extensive distribution from the open tundra to moorlands and subarctic forests at lower altitudes. Willow ptarmigan, with its white winter plumage, is adapted to harsh winter conditions, and it burrows in snow as protection from cold weather and predation (Andreev, 1991; Hoglund, 1980). Willow ptarmigan exhibits large annual fluctuations attributed to variation in breeding success (Myrberget, 1974; Steen and Erikstad, 1996). Adult mortality, nest loss, and chick mortality have mostly been attributed to predation (Erikstad et al., 1982; Marcström and Höglund, 1980; Smith and Willebrand, 1999; Steen et al., 1988). Population change show negative density–dependent growth (Pedersen et al., 2004), but the variation in breeding success is not related to adult abundance (Hörnell-Willebrand et al., 2006).

A recent review of the long-term population trends of willow ptarmigan identified negative trends in parts of its distribution (Fuglei et al., 2020), and several potential factors could affect adult survival and breeding success negatively (Henden et al., 2017). The ongoing changes in climate is expected to result in winters with shorter snow seasons, which could increase the vulnerability of the white ptarmigan on snow-free ground to birds of prey (Henden et al., 2020; Melin et al., 2020). On the other hand, air temperature after snow-melt is important for the timing of spring, with warmer conditions contributing to earlier phenology in vegetation communities (Kelsey et al., 2021). Early onset of spring with appearance of new nutritious vegetation positively affects nutritional status and breeding conditions (Brittas, 1988; Eriksen et al., 2023; Erikstad and Andersen, 1983; Marcström and Höglund, 1980).

Historically, willow ptarmigan has been an important game species and commodity in game markets (Hjeljord and Loe, 2022; Lloyd, 1867; Sotherton et al., 2009). Today, its value as a game for consumption has declined, and it is mostly a species of interest for sport hunting. However, the need for harvest management has been addressed in recent studies (Aanes et al., 2002; Pedersen et al., 2004; Smith and Willebrand, 1999). In response to these concerns, national schemes of line transects with pointing dogs and distance sampling (Thomas et al., 2010) have been developed to estimate densities in Sweden, Norway and Finland the last 15 - 25 years (Eriksson et al., 2006; Helle et al., 2016; Nilsen et al., 2024). Systematically collected field data over several decades provide important insights in both short- and long-term dynamics of ecological systems (Lindenmayer et al., 2012; Magurran et al., 2010). Time series of harvest statistics and catch data have provided valuable insights into population dynamics (Keith, 1963; Royama, 1992). However, long-term studies on counts and estimates of abundance are less common among wildlife species, but see (Jacobson et al., 2004; Ranta et al., 1995; Sæther et al., 2008).

The recent development of hierarchical state-space models in a Bayesian framework has improved the possibility of jointly analyzing several different data sources of wildlife time series and increased the possibility of identifying factors that are important for population change (de Valpine and Hastings, 2002; Kéry and Royle, 2016; Lee et al., 2015). A great advantage is that they can provide separate estimates of observation and process error, which otherwise tend to bias estimates of model parameters (Aanes et al., 2002; Freckleton et al., 2006). These models are well suited to iteratively using new data, updating short-term predictions, and supporting management decisions (Bowler et al., 2020; Frost et al., 2022; Henden et al., 2020). The main aim of this study was to investigate the drivers of short-term population dynamics of the adult willow grouse and assess potential population trends over the last 50 years. To achieve this we used a unique set of line-transects from the Lövhögen area in Sweden collected between 1963 and 2023 and developed a hierarchical state-space model to estimate changes in adult population density and breeding success. We estimated the effects of vole abundance and spring conditions on breeding success, and their effect on adult population change. The lack of snow in autumn and spring was also used as predictor of adult change. The model includes parameters for density dependence and a linear trend for adult change.

## Material and methods

### Study Area

The study area, Lövhögen 80 *km*^2^, is located in the southern part of (62’N, 13’E) of the Swedish alpine tundra, where the landscape is gradually transformed into continuous boreal conifer forest. The altitude varies between 750 and 1 000 m.a.s.l. and is dominated by birch forest and open tundra (Brittas, 1988). The tree line is about 800 m.a.s.l., and about 20% is wooded mainly with mountain birch *Betula* spp.. There are small scattered stands of Norway spruce *Picea abies* and Scots pine *Pinus silvestris* above the tree line and a few scattered willow *Salix spp*. thickets. Almost one-third of the area is peatland, characterized by sedge *Carex* spp., cloudberry *Rubus chamaemorus* and cotton grass *Eriophorum* spp. About half of the study area is heathland, dominated by dwarf shrubs such as heather *Calluna vulgaris*, bilberry *Vaccinium myrtillus*, crow berry *Empetrum hermaphroditum* and dwarf-birch *Betula nana*. The climate is continental, with temperatures averaging -10 C° in January and snow cover stay for about 6 months in winter. The average temperature is +14 C° for July. There are few lakes, but frequent small creeks drain the area into larger streams. The study area is privately owned by a single landowner, and the harvest of willow grouse is highly restricted (Aanes et al., 2002).

### Data

#### Line transect survey

A permanent monitoring program based on line transect surveys using pointing dogs was established in 1963 to estimate the abundance of adult willow ptarmigan. A total of 18 lines, 600 m apart, with a total distance of 139 km cover the entire study area. The start and end points of each line were marked with permanent poles. These lines have remained unchanged since the survey began. The perpendicular distance to the observed willow ptarmigans was used to estimate the probability of detection. Initially, the effective strip width (ESW) was calculated as the average of the observed distances within 200 m of the line. In the period A between 1963 and 2004, counts were made in late June. The perpendicular distance to all detected adults were registered and, sex was determined by plumage and/or behavior. It was not possible to determine the sex of some adults, and the observed ratio of males to females was used to estimate the number of undetermined sex. In late June, females are usually incubating, and are therefore very difficult to detect. Therefore, the estimated number of adult males was assumed to correspond to a pair of two adults. Unfortunately, the distances between 1963 and 1975 were destroyed when the Boda wildlife research station burned down, but the estimated adult densities for that period are still available. The distances were lost in 1979, 1980, 1985 and 1986, and registered as missing values in the data. The model development was restricted to the period with available distances that starting in 1976. The original estimates of adult densities between 1963 and 1975 are presented in the appendix. In period B from 2004, the study area was included in the national monitoring program for willow ptarmigan in the tundra (Eriksson et al., 2006), and the counting was delayed to late July or early August, similar to other areas. In period B, it was possible to observe both adult and brood size in the field. It was not assumed that a brood was always accompanied by a pair because sex segregation among adults may occur already in August (Marcström and Höglund, 1980). The number of adult observations per year varied between 73 and 206, with a median of 129 during the study period. The survey was completed within a week, and most members of the monitoring team have remained for many years.

### Brood size

Between 1963 and 1994, brood size was estimated from the age ratio of harvested willow ptarmigan. Harvest management in Lövhögen is aimed to sample the population and differs from many other areas. Hunting occur during 5 - 7 days after the season opens 25th of August, and a previous estimate of harvest rate was on average 0.144 but with large variation (s.d. = 0.106) (Aanes et al., 2002). Hunters are divided into 2-4 teams with several hunters and one professional dog handler. The area is divided into sections that are visited only once to ensure that the effort is equal in all parts of the area. In 1963 and 1971, the relationship between the age distribution of flushed willow ptarmigan in August was compared to the age ratio of harvested willow ptarmigan in early September. The difference in age ration was minor, 66% and 74% in counts and 63% and 70% in harvested birds (Marcström and Höglund, 1980). From 2004, when counts were made in early August, we used the number of observed chicks per two adults as the average brood size.

### Vole numbers

Voles were snap-trapped in late June and later in early August during in the same time as the willow ptarmigan surveys, and they accumulated between 200 and 600 trap nights each year. The logarithm-transformed number of voles per 100 trap nights was used as a vole index (Hörnfeldt, 1978), data were standardized for each period and then combined. Three years lacked observations (1976, 1977 and 1990). The first initial two years of the study were replaced by the median of the complete time series. It was possible to replace the third missing observation in the series with the average for 1989 and 1991.

### Meteorological data

Data on snow depth was obtained from the Swedish Meteorological and Hydrological Institute for the Särna station (61.70, 13.13), 437 m.a.s.l. The station is about 28 km southwest of Lövhögen. Data were not available for May in 2000 due to the renovation of the station; therefore, we used the average number of observations for each time and day in May in 1999 and 2001 as data for 2000.

The number of snow-free days was extracted for spring (April and May) and autumn (October and November). The number of snow-free days was standardized and included as a covariate of adult change. We extracted the ambient temperatures for all days in May at 03:00 h during the study. The last day with a temperature less than -2.2*C*° has been validated as a measure of spring vegetation growth (Schwartz et al., 2006). We used as an index of annual variation in availability of newly-growing and nutrient-rich plants. The dates were standardized and included as covariate in the model for breeding success.

**Table 1.** – Overview of the parameters estimated in the hierarchical models of adult density and breeding success. The breeding success model provide input to the model of adult change. For more information refer to the analysis section.

| Model | Parameter |
| --- | --- |
| Population dynamics | Breeding success the previous year<br>Snow-free days in spring the same year<br>Snow-free days in autumn the previous year<br>Density dependence<br>Linear trend |
| Breeding success | Vole abundance<br>Late frost |

### Analysis

We developed a hierarchical state-space model in JAGS (Plummer et al., 2003) following (Henden et al., 2020; Kéry and Royle, 2016). We used Markov Chain Monte Carlo (MCMC) methods to estimate the posterior distribution of parameters and adult density estimates. The model was run for 30 000 iterations with a burn-in of 8 000. The thinning was set to 1, and three chains were used to evaluate convergence. The package MCMCvis (Youngflesh, 2018) was used to evaluate the trace plots and posterior distribution of the model parameters. All parameters had an R-hat value less than 1.01. The main model and submodel for breeding success were validated by simulating replicate data and comparing predictions with both original and simulated data (Schaub and Kéry, 2021). The Bayesian P-value was close to 0.5 for both the main and breeding success model (0.49 and 0.41 respectively). Parameter estimates are presented with a 95% credible interval (Cr. I), and the plot of the estimated time series of adult densities show the 80% highest posterior density. All analyses and graphics were performed in R (R Core Team, 2024) and the JagsUI package was used as an interface to JAGS from R (Kellner, 2024).

#### Distance sampling and the probabilityof detection

Distance data were prepared for analysis using a cutoff of 200 m and a bin width of 20 m. This resulted in 10 distance intervals, where each observation was categorized into a specific distance interval (1-10). During the 48 years in which distances were available we included 3531 distances less than 200 m. Thus, the line transects covered an area of approximately 55 *km*^2^.

The detection probability was computed in a multinomial model of cell probabilities, assuming a half-normal detection function. See eq. 1 where *x* denotes the midpoint distance of the distance class. All distances were pooled to estimate the detection probability, and we did not add any covariates to the detection equation. However, we included separate intercepts of the estimate of *log* (*σ*_*A,B*_) to evaluate the potential effect of the different periods on the probability of detection (*pcap*_*A,B*_). Further information can be found in the model code in the Appendix, and in chapter 9.7 in Kéry and Royle (2016).

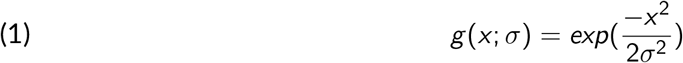

#### Model of breeding success

We estimated the effects of vole abundance and spring phenology on standardized brood size and used these estimates as an independent variable in the part of the model that describes adult changes. This made it possible to estimate breeding success for the period 1995 - 2003 when age ratio of harvested willow ptarmigan was not available due to changes in harvest management. We used standardized data on average brood size, vole abundance and late frost in May to estimate annual changes in breeding success. Data on brood size were positively correlated with the vole index (0.324 (P = 0.044) but negatively correlated with late frost (−0.322, P = 0.045). However, the vole index and late frost were not significantly correlated (−0.132, P = 0.369). We assumed that the error terms were normally distributed, (eq. 2).Steen et al. (1988) present previous analysis on the effects of vole abundance and weather on breeding success in our study area.

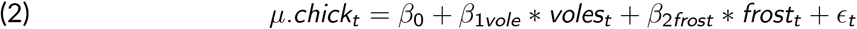

### Main model of adult change

We used two process models: one without covariates for the first year and a second with covariates for subsequent years. In the section without covariates, the initial density and observation error (*µ*_*t*_ = *β*0_*A*_) are estimated (Sillett et al., 2012). The second part include covariates and models population change in subsequent years (eq. 8) including process variation (eq. 6, *PrOc*). The number of observations is assumed to follow a binomial distribution based on the probability of observation (pcap) and the number of individuals (N) (eq. 3), which are assumed to show a Poisson distribution with mean *λ* (eq. 4). *λ* is the product of density and the area covered by the line-transect surveys (eq. 5). The logarithm of the expected density (*µ*_*t*_) is assumed to be normally distributed (eq. 6). We then use a Gompertz-model to estimate the change in adult densities (*log* (*D*_*t*_)) in subsequent years. *β*0_*A,B*_ is the intercept for the two different sampling periods, *β*1 is the strength of density dependent and subtracted with 1 to scale the parameter such that 0 is lack of density dependence. *β*2 is the parameter of the linear trend, *β*3 is the effect of variation in estimated breeding success, *β*4 is the effect of snow-free days in spring in the same year, and *β*5 is the effect of snow-free days in autumn the previous year.

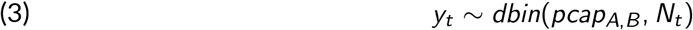

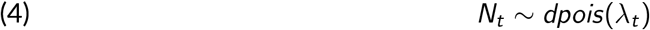

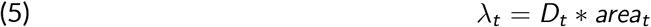

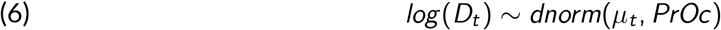

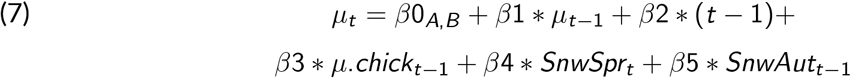

Following Royama (1992) we compared the first and last 20-years of the estimated adult densities. We compared the mean (t-test) and variance (F-test) of the two periods, and examined the autocorrelation plots for significant coefficients using the Bartlett band.

## Results

### Adult dynamics

Adult density was on average 5.4 adults per *km*^−2^, with an interquartile range of 4.9 to 6.4 (Fig. 1a). The coefficient (*β*2) for the linear trend of logarithmic adult density was small and overlapped with zero, (0.007, 95%Cr.I. -0.007 to 0.021) (Fig. 2a). The mean logarithmic density for the first and second 20th year periods was 1.773 and 1.649 respectively (t = 1.700, df = 37.284, P = 0.097), and similar ratio of variance (F = 0.584, numerator df = 19, denominator df = 20, P = 0.246). In addition, the autocorrelation plots did not show any significant coefficient at time lags 1,2, …10, indicating a stationary process.

**Figure 1.**
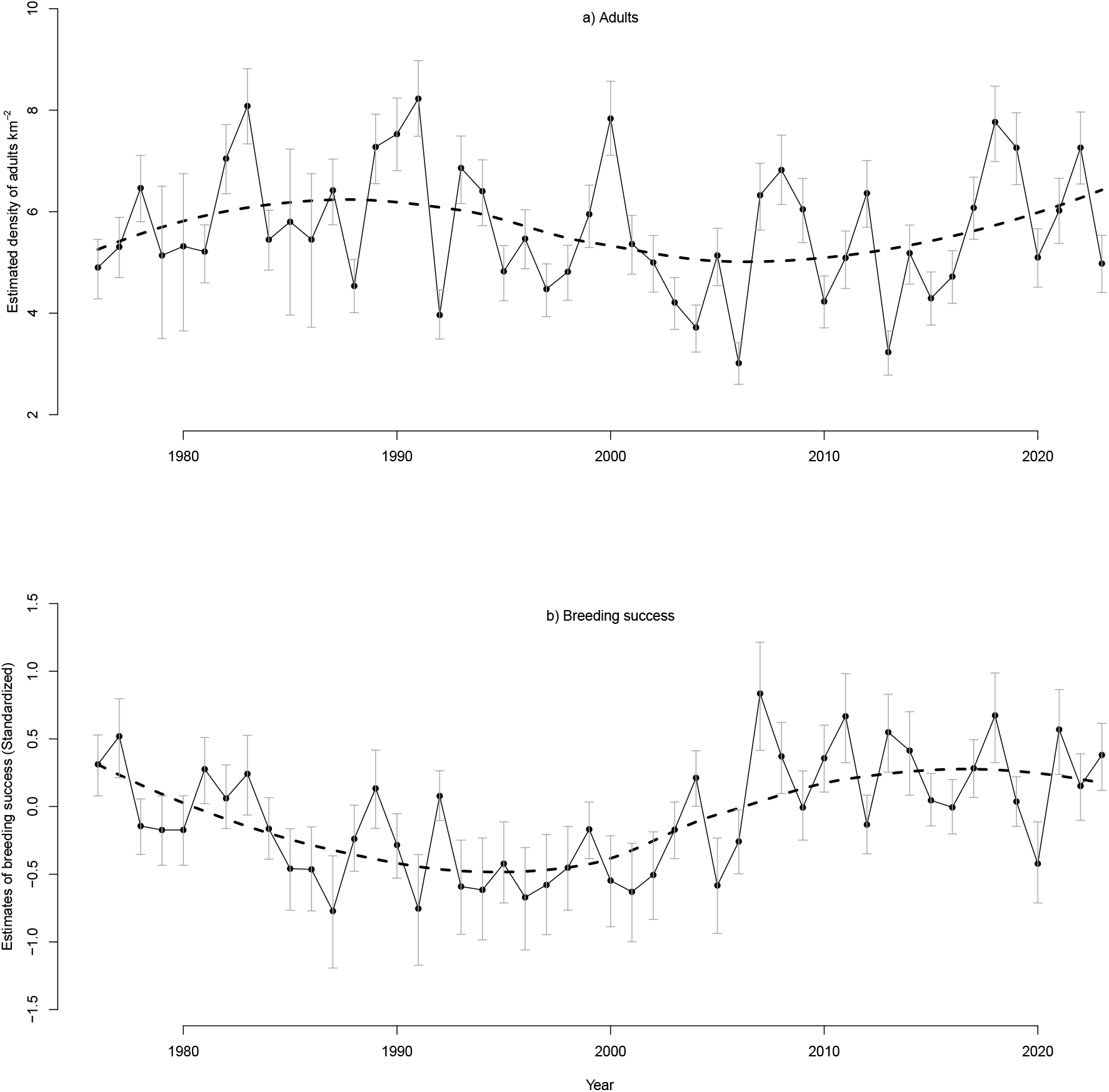
The estimated dynamics of a) adult densities and b) breeding success during 1976 - 2023. The vertical lines show the 80% highest posterior density interval. The smooth stippled lines are produced by the loess function.

**Figure 2.**
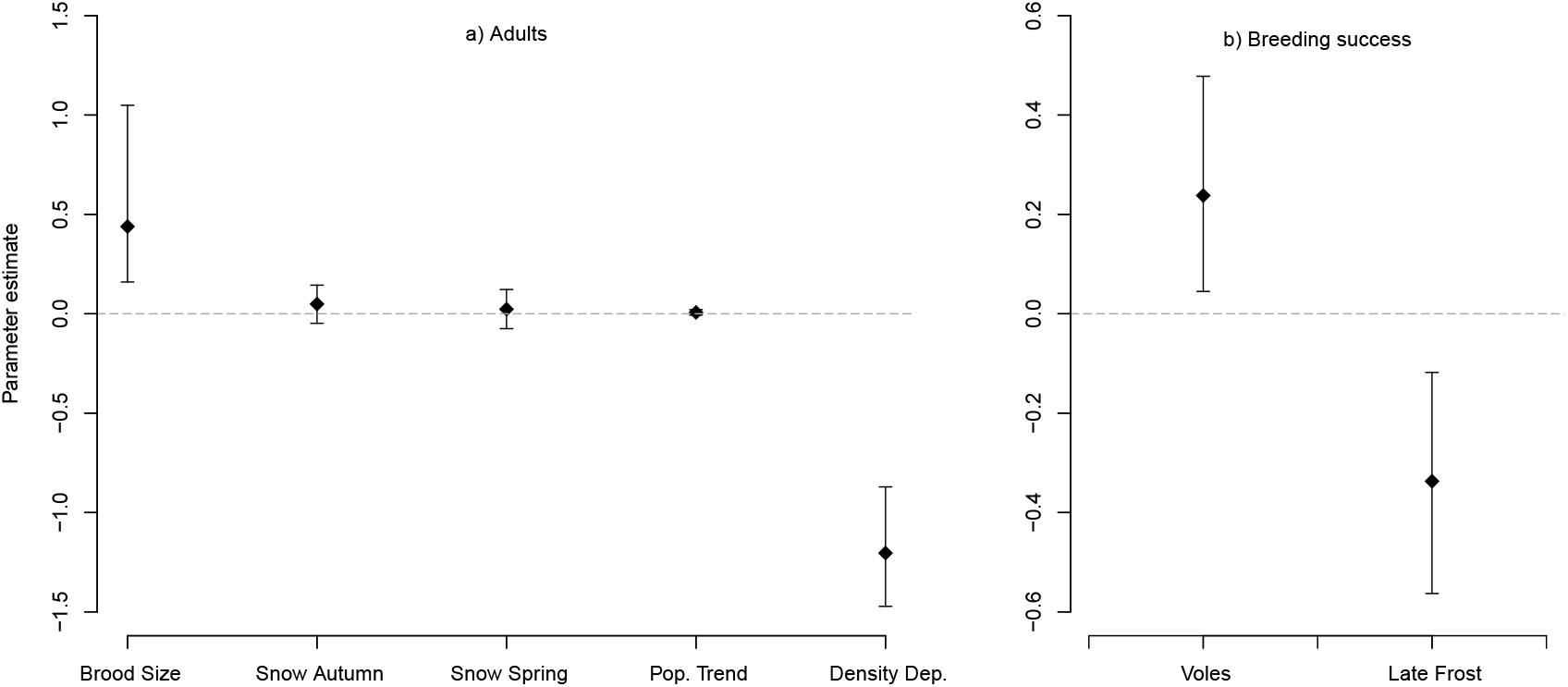
Parameter values and 95% credible intervals for the explanatory variables of annual change in the a) adult density and b) breeding success.

The parameter for direct density dependence was -1.204 (*95%Cr*.*I*., -1.472 to -0.871), which indicates a strong negative density dependence. The parameter for estimated breeding success was clearly positive, 0.439 (*95%Cr*.*I*., 0.160 to 1.049). The parameters for snow free days in the previous autumn and current spring was positive but non-significant, 0.049 (*95%Cr*.*I*., -0.048 to 0.144) and 0.023 (*95%Cr*.*I*., -0.075 to 0.122), respectively. See figure 2a.

The errors for the first year (sdperctau) and subsequent years (sdproctau) were estimated to 0.595 (*95%Cr*.*I*. 0.077 to 0.980) and 0.212 (*95%Cr*.*I*., 0.159 to 0.280) respectively. The intercepts for periods A and B of the second process model were 2.126 and 1.611, indicating a higher adult density in the first period but the 95% credible intervals greatly overlapped (1.551 - 2.675 and 0.966 - 2.359 respectively).

### Breeding success, voles and late frost in May

Breeding was positively affected by vole abundance, (0.238 with *95%Cr*.*I*., 0.045 to 0.478), but negatively affected by late frost in May (−0.337 with *95%Cr*.*I*., -0.563 to -0.118; Fig. 2b). The adult density estimates did not correlate with breeding success, (−0.060, t = -0.409, df = 46, P = 0.684). The coefficients of the partial autocorrelation function at lags 1 and 3 were positive and 95% confidence intervals (Conf.I.) did not overlap zero, 0.426 (95%Conf.I. 0.143-0.709) and 0.382 (95%Conf.I. 0.099-0.283), respectively.

### Changes in snow-free days and last frost in May

There was a large variation in the number of snow-free days in autumn and spring, and a significant linear increase trend over time, (slope=0.040 *R*^2^ = 0.274, Std. Err. = 0.006, t= 6.093, P<0.0001). See figure 3a. From 1976 to about 1995, the last frost in May tended to occur increasingly later, followed by a period when the date of the last frost decreased to earlier dates again (Fig. 3b). The values of last frost in May was not correlated with the snow-free days in spring or autumn (r<0.011, P>0.94).

**Figure 3.**
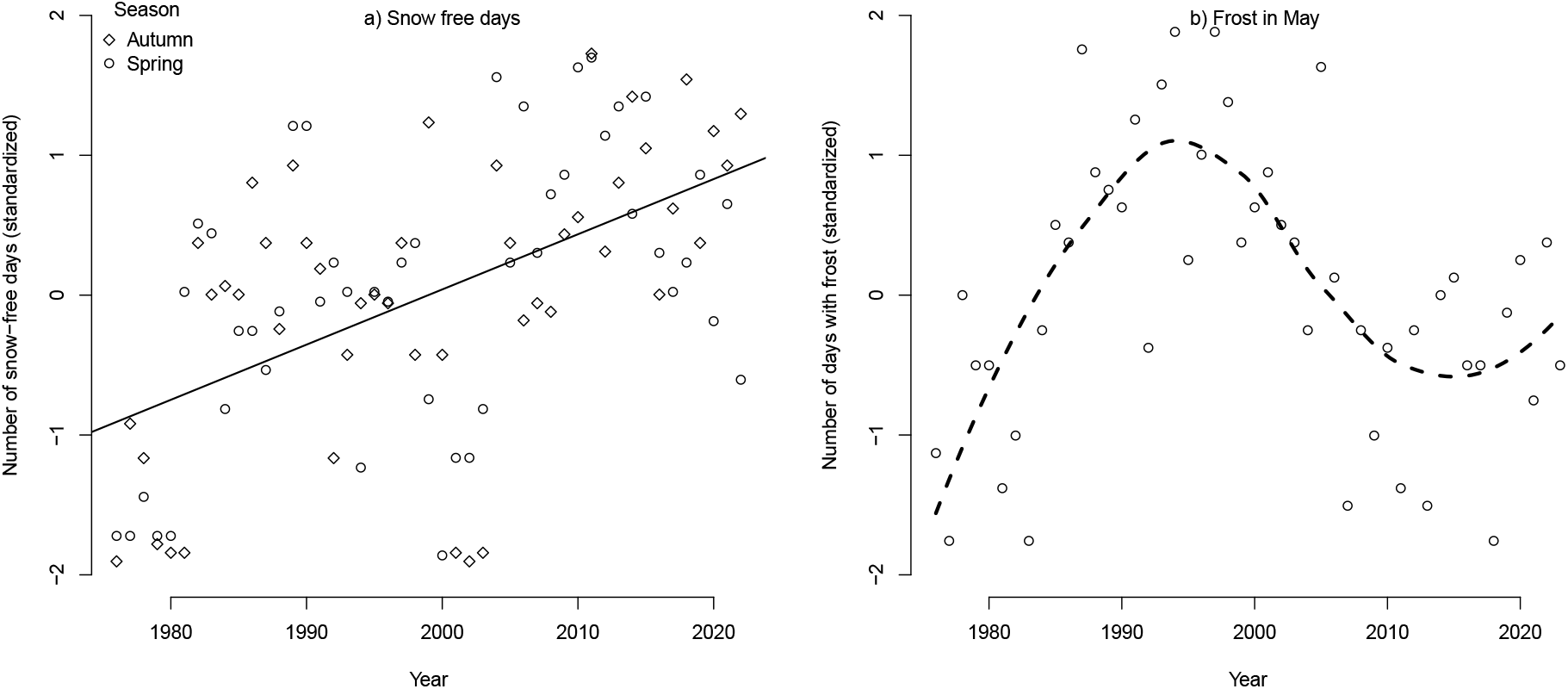
a) The increase in snow-free days in autumn (October and November) and subsequent spring (April and May). The regression slope is positive and statistically significant, 0.040 (Std. Err. = 0.006 t= 6.093, P<0.0001). b) Last day in May with temperature below -2.2*C*°. The smooth stippled lines are produced by the loess function.

### Detectability

The probability of detecting willow ptarmigan within the *±*200 m strip around the line transect in period A and B was 0.441 (*95% Cr*.*I*., 0.423 - 0.459) and 0.432 (*95% Cr*.*I*., 0.412 - 0.453) respectively. The average of these two estimates (0.436) corresponds to an effective strip width of 87.2 m, which is substantially longer than the 63.8 m that was used previously in the period 1963 - 1975. (See figure 4 in appendix.)

**Figure 4.**
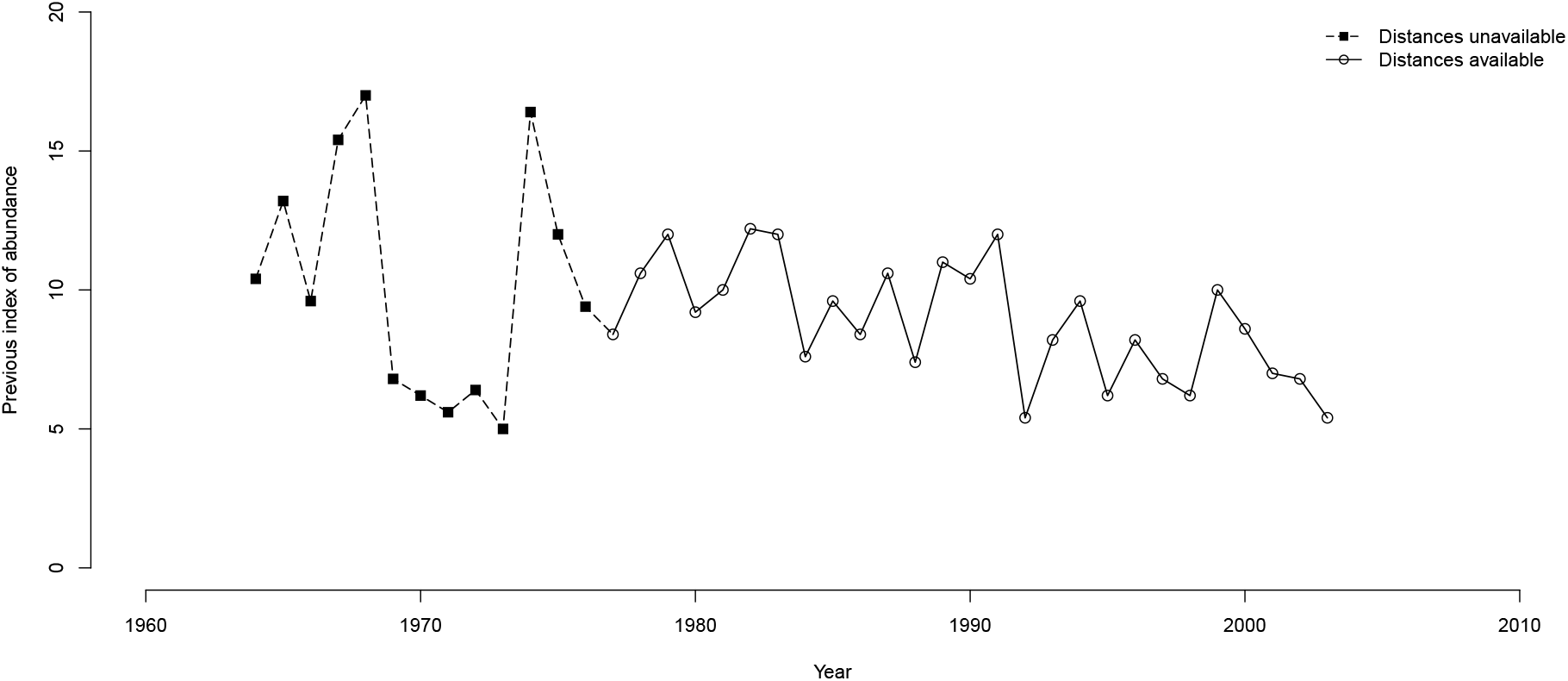
The original time series based on a strip width estimate of twice the average distances recorded. Note that distance measurements between 1963 to 1975 was lost in a fire.

## Discussion

The estimates of adult density did not show any trend for the period 1976 to 2023, and the dynamics were characterized a strong direct negative density dependence. We propose that the functional response by the predator guild in the region is the most likely process that explains these results. In addition, large-scale natal dispersal and recruitment of juvenile females can also contribute to stabilizing the population (Hörnell-Willebrand et al., 2014; Pulliam and Danielson, 1991). Earlier studies have shown that predation on young and adult willow ptarmigan is high during autumn, both in areas open and closed to harvest (Eriksen et al., 2025; Israelsen et al., 2020; Smith and Willebrand, 1999). However, our results do not support the hypothesis that lack of snow in autumn and spring increases vulnerability to predation and population decline, as suggested by Melin et al. (2020). In fact, recent studies have shown that in northern areas, more precipitation during winter with still below freezing temperatures leads to more snow in early spring, which might counteract earlier snowmelt from increase spring temperature (Bjerke et al., 2025). While this is one of the longest existing time series on willow ptarmigan density in Scandinavia, there is some uncertainties about the trend assessment since 1) the timing of the yearly monitoring was change during the course of the study and 2) the initial years of this time series (1963-1975) could not be included in the distance sampling model due to missing data, while this period appears to show the most extreme fluctuations and the highest adult estimates in the original time series. In fact a population decline during the late 60’s with short-term fluctuations at lower densities seems to be in line with the population decline in willow ptarmigan in southeastern Norway during the last 150 years (Hjeljord and Loe, 2022). Similar declines and changed fluctuation patterns were reported in capercaillie (*Tetrao urogallus* and black grouse *Lyrurus tetrix* during the 60’s (Jahren et al., 2016). Further, the climate data (snow and temperature) were retrieved from the weather station closest to the study area, which was located at a considerably lower elevation (437 m.a.s.l compared to the ptarmigan monitoring at approximately 750 – 1100 m.a.s.l), potentially representing different climatic conditions relative to the study area. Despite these challenges, we believe this analysis provide valuable insight from a unique long term time-series about the population dynamics of willow ptarmigan, and supports many of the conclusions on willow ptarmigan biology from earlier studies.

Breeding success is an important driver of changes in, but independent of, adult density. Our findings agree with numerous previous studies that have concluded that predation on nests and chicks follows fluctuations in voles and lemmings (Steen and Haugvold, 2009; Steen et al., 1988). Willow ptarmigan is well adapted to winters with deep snow, and the birch forest provides an ample source of twigs and catkins (Brittas, 1988; Hoglund, 1980). Small herbivores in northern areas feed regularly, emphasizing energy conservation rather than acquiring energy reserves (Thomas, 1987). Extremely cold weather and high precipitation during the first days after hatching can be detrimental (Bowler et al., 2020; Erikstad and Spidsø, 1982; Marcström and Höglund, 1980), but Steen et al. (1988) showed that the variation in weather after hatching was poorly correlated with breeding success. In stead, we propose that climate conditions during egg formation and laying are more important. Females generally form eggs when plants begin to grow in spring (Moss, 1997), and late spring is associated with delayed egg laying (Wiebe and Martin, 1995). In our study area, the variation in hatching date was on average 12 days (17 - 29 June) (Marcström and Höglund, 1980), and Brittas (1988) showed that the digestibility of spring food in willow ptarmigan crops was positively correlated with both female body condition during egg laying and subsequent breeding success. This is in accordance with the maternal hypothesis, which states that female body condition during egg laying and incubation affects the survival of chicks and subsequent recruitment (Blomqvist et al., 1997; Coslovsky and Richner, 2011). The negative effect of late frost in May reduces the availability of newly-growing and nutrient-rich plants, such as shoots of cotton grass *Eriophorum* spp. We believe it is important to explore the potential for vole abundance in early summer and the date for the last frost in May to provide an early indication of the average brood size in August.

Despite large annual variations, the long-term average onset of spring began earlier in our study area (Jin et al., 2019). However, we did not observe any effect of the increase in snow-free days in autumn and spring, but there are a number of possible mechanisms by which a warmer climate could affect willow ptarmigan population dynamics. For example, short periods of rain in winter can form an icy crust on both snow and birch twigs, which may force willow ptarmigans to leave an area (Hoglund, 1980). Another mechanism would be changes in the vegetation community (Bråthen et al., 2024; Tuomi et al., 2024).

The characteristics of the willow ptarmigan population dynamics are likely to vary depending on the food-web structure. The structure of trophic levels becomes simpler in the north compared to further south (Callaghan et al., 2004b), where the tundra becomes fragmented and surrounded by continuous boreal forests. At lower latitudes, animal diversity increases with various herbivores and predators (Callaghan et al., 2004a). Patterns of population dynamics depend on trophic interactions and time lags (Turchin and Taylor, 1992), and we expect willow ptarmigan populations to show increased densities and amplitudes along a geographical gradient from south to north similar to vole populations in Scandinavia (Erlinge, 1987; Soininen et al., 2025; Turchin and Hanski, 1997) and snowshoe hares (*Lepus americanus*) in North America (Krebs, 2010). Our study area is in the southern range of the tundra, and we suggest that a diverse predator assemblage makes it difficult for the population to escape a top-down control. Further north, with fewer predator species, it may be more common to escape top-down control that might result in higher amplitudes and densities of willow ptarmigan populations.

## Acknowledgements

We are grateful to Stora-Enso for allowing us to perform fieldwork and provide accommodation in the field. Their support has been fundamental during this long-term study. We thank the counting crew who volunteered with their highly qualified pointing dogs over the years. Their commitment and understanding of the required precision of data cannot be underestimated. We thank the reviewers of the previous versions of the manuscript for helpful comments and suggestions. A preprint version of this article has been peer-reviewed and recommended by PCI Ecology (https://doi.org/10.24072/pci.ecology.100540).

## Conflict of interest disclosure

The authors declare that they comply with the PCI rule of having no financial conflicts of interest related to the content of the article.

### Original estimates of adult densities, 1963 - 2003

#### Script and codes

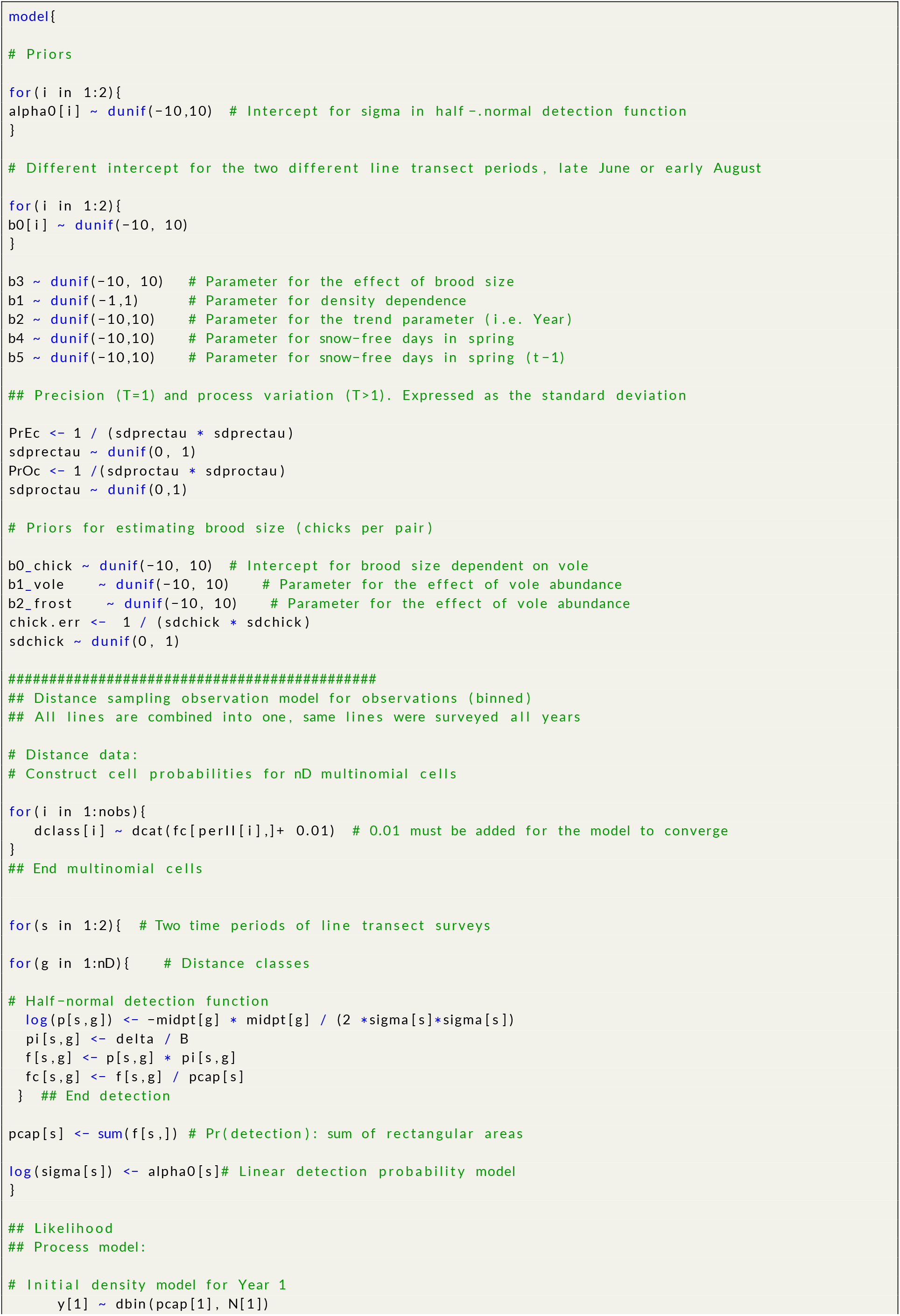
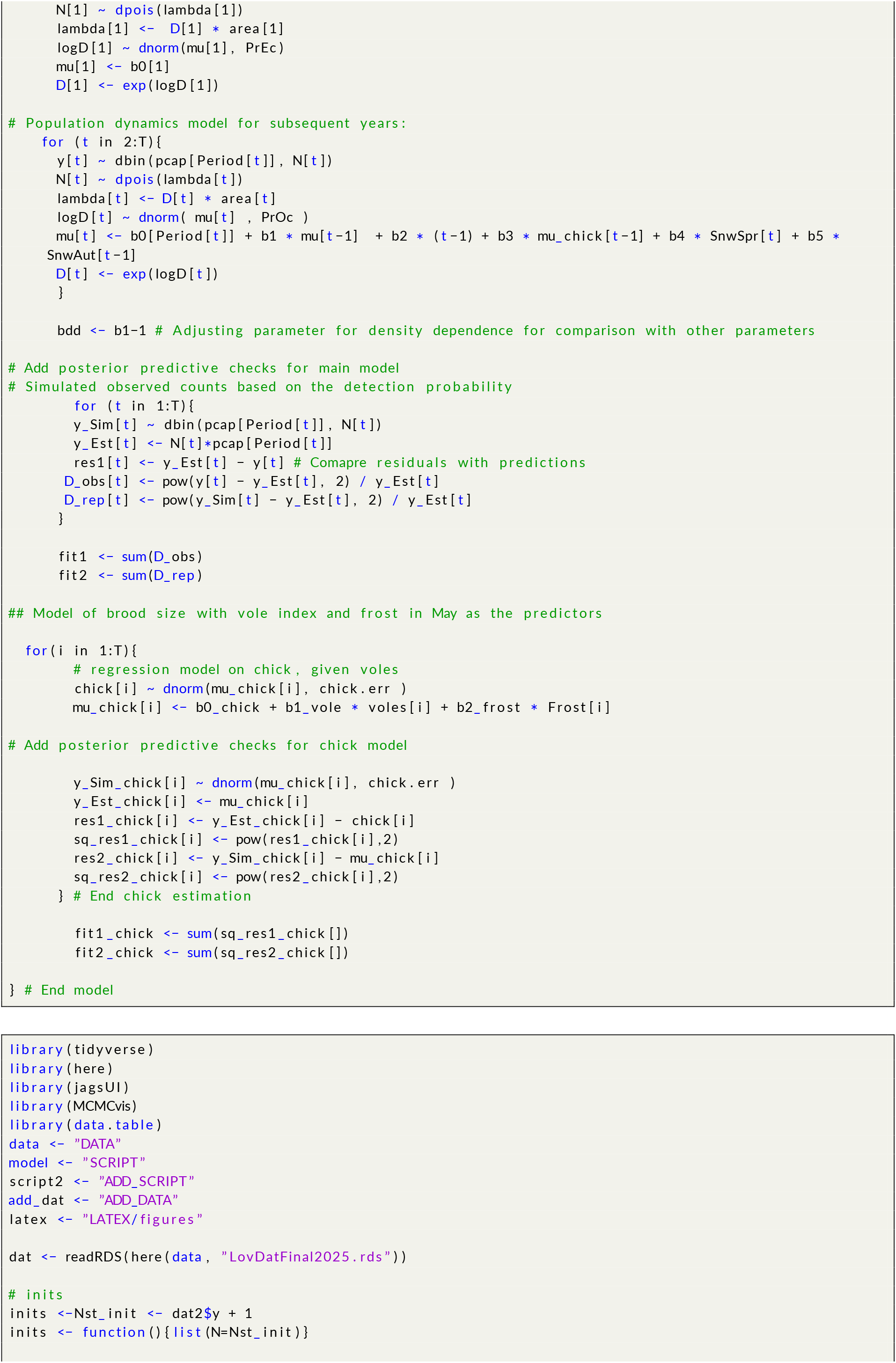
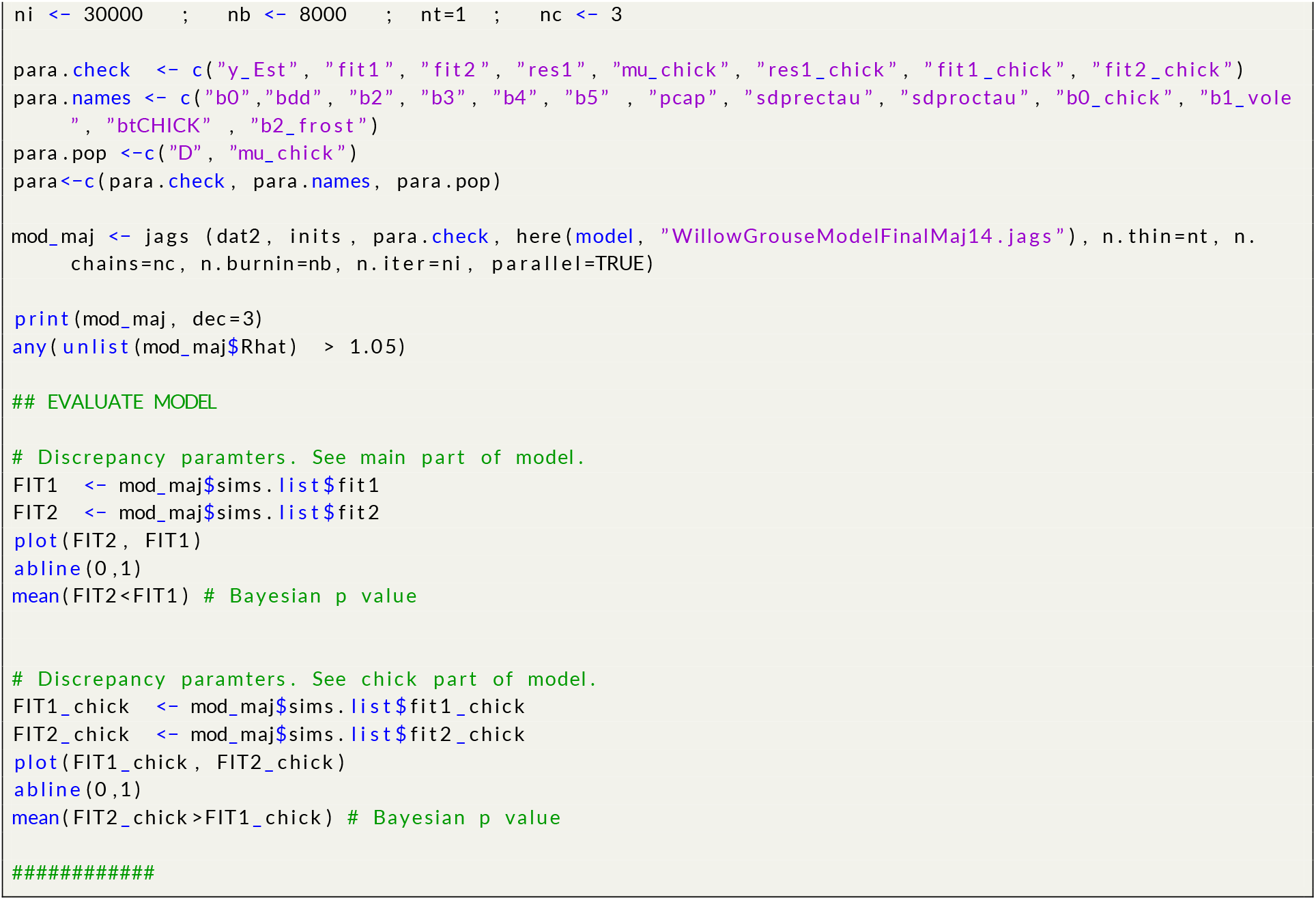

Data and scripts are available online.

